# Exploring causal factors of coastal chlorophyll-a dynamics and their potential contributions to near future forecasting

**DOI:** 10.64898/2026.01.07.698127

**Authors:** Suixuan Huang, Masayuki Ushio

## Abstract

Harmful algal blooms have been causing significant damage worldwide, and Hong Kong is no exception. To understand the drivers of algal bloom formation and forecast the dynamics of chlorophyll-a (Chl-a), a proxy for algal abundance, in Hong Kong waters, this study utilized nonlinear time series analysis, called empirical dynamic modeling (EDM), to investigate Chl-a dynamics using *in situ* measurements and remote sensing data. We first conducted causality tests of EDM to identify environmental factors influencing Chl-a at different sites. As for the *in situ* measurement data, salinity was the strongest causal factor among environmental factors. However, inputting the causal factors into the forecasting model did not greatly improve the forecasting performance for Chl-a, suggesting that factors not included in the current dataset, such as wind direction and current speed, may play a more critical role in Chl-a dynamics. As for the remote sensing data, sea surface temperature (SST) showed significant causal effect on Chl-a at most sites and the multivariate forecasting model including Chl-a and SST outperformed the univariate model at most sites. This study is the first to employ EDM to investigate Chl-a dynamics in Hong Kong waters, showcasing its potential to identify causal factors and improve forecasting accuracy. The findings provide scientific insights into Chl-a dynamics and water quality monitoring and modeling in a coastal region.

## 1. Introduction

Harmful algal blooms (HABs), commonly known as “red tides,” refer to the discoloration of coastal waters caused by the rapid growth and accumulation of harmful microscopic algae (phytoplankton) (Zohdi & Abbaspour, 2019). These blooms can produce toxins triggering foodborne illnesses, such as amnesic shellfish poisoning and ciguatera fish poisoning (Lopes et al., 2019; Pradhan et al., 2022; Wang, 2008). Zooplankton, filter-feeding shellfish and herbivorous fish consume these phytoplankton, acting as mediators for toxin transfer within the food web, ultimately affecting humans. Even non-toxic blooms can indirectly harm marine life by depleting oxygen during decomposition (Flewelling et al., 2005). Due to the great impact on public health concerns, the seafood industry and tourism, the economic loss is considerable. In Florida, for instance, red tides have been observed since the 1840s and the annual losses reach millions of dollars (Kirkpatrick et al., 2004). To prevent such damage caused by HABs in coastal ecosystems, it is essential to understand the mechanism and predict the outbreaks of red tides.

The concentration of chlorophyll-a (Chl-a) is a common proxy for assessing phytoplankton abundance. Thus, identifying the drivers of Chl-a dynamics contributes not only to developing effective HAB monitoring plans but also to predicting HAB dynamics. Previous attempts to identify drivers of the dynamics of Chl-a or phytoplankton community composition mainly relied on correlation-based methods, for example, linear regressions, Canonical-Correlation Analysis (CCA), and Redundancy Analysis (RDA) (Deconinck et al., 2025; Li et al., 2023; Yin, 2003). These methods assume that environmental variables are independent and the effects of such variables are separable from each other, which are features of linear systems. However, the effectiveness of these methods could be limited when studying nonlinear systems such as marine ecosystems, where variables are interdependent upon each other (Glaser et al., 2014). Moreover, correlation may misidentify causal variables, as “correlation does not imply causation” (Berkeley, 1988). That is, correlation can occur without causation, and causation may also occur in the absence of correlation.

As an alternative non-parametric method, Empirical Dynamic Modeling (EDM) was developed to analyze complex dynamics found in natural ecosystems (Anderson et al., 2021; Glaser et al., 2014; Sugihara et al., 2012; Sugihara & May, 1990). Instead of deriving equations to describe how the variables are related or how the system evolves, EDM delineates a “trajectory” of the system evolution in a high-dimensional state space (i.e., a manifold) (for example, see figures in Ye et al., 2015). EDM was originally developed to make near-future forecasts for deterministic complex systems in the state space built by lagged time series of a single variable (Sugihara & May, 1990). Eventually, several tools of EDM were developed to analyze the relationships among multiple variables within the same system (in HAB studies, it would be time series of Chl-a and environmental factors), enabling detection of causalities among variables (Sugihara et al., 2012) and quantification of interaction strengths (Deyle et al., 2016), and EDM tools have been applied to various ecological time series to understand and forecast ecosystem dynamics (Deyle et al., 2022; Tsai et al., 2024; Ushio, 2022; Ushio et al., 2018).

Hong Kong is a coastal city with a long historical record of HABs. Located to the east of the Pearl River Estuary (PRE) and to the north of the South China Sea (Fig.1a), Hong Kong is affected by fresh water and sea water at the same time (Wong et al., 2007). As a result, there are significant spatial and seasonal variations in water circulation, stratification, salinity, temperature, and nutrient levels there. In this study, we aim to detect causal factors of Chl-a dynamics in Hong Kong waters and improve the near-future forecasting performance using EDM tools. Our work is built on the long-term *in situ* observation of Chl-a and environmental factors in 76 sites (Fig.1a) conducted by the Environmental Protection Department (EPD) of the Hong Kong government. Additionally, remote sensing measurements collected by sensors such as Sea-Viewing Wide Field-of-View Sensor (SeaWiFS) and Moderate Resolution Imaging Spectroradiometers (MODIS) provide great support for monitoring Chl-a and other parameters with high coverage and fine spatial resolution (Chen et al., 2013). In Hong Kong, MODIS data at 4-kilometer resolution include 176 sites in total, of which 101 are located on the water area (Fig.1b). The spatial resolution of remote sensing data is also fine, allowing for daily or weekly observations. However, frequent occurrence of missing data in daily or finer temporal resolution data complicates the understanding of spatiotemporal variations (Zhang et al., 2025). Monthly-averaged data may mitigate the effect of missing data, but the reliability of remote sensing data in coastal regions such as Hong Kong waters—where particle suspension significantly impacts remote sensing images—has not been thoroughly investigated for Chl-a monitoring, although the monthly remote sensing data is generally well correlated with the *in situ* measurement data (Fig. 1c). Comparing the *in situ* measurement data and remote sensing data would give us an opportunity to test the reliability of the remote sensing data for Chl-a monitoring in coastal regions.

**Figure 1.**
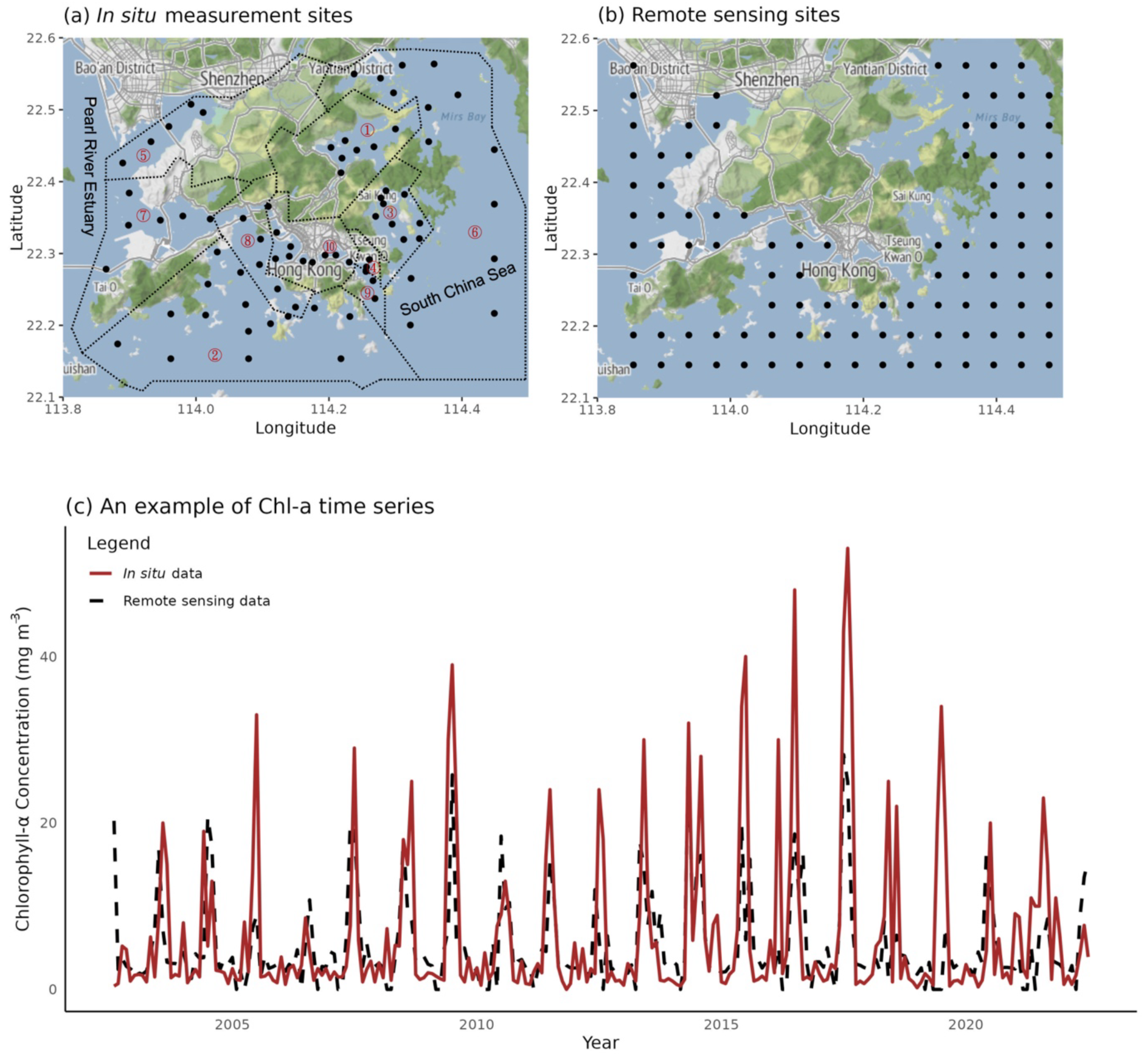
Study sites of (a) *in situ* measurements conducted by EPD and (b) remote sensing collected by MODIS, represented by black points. In (a), there are ten water control zones declared by EPD, namely ①Tolo Harbour and Channel, ②Southern, ③Port Shelter, ④Junk Bay, ⑤Deep Bay, ⑥Mirs Bay, ⑦North Western, ⑧Western, ⑨Eastern and ⑩Victoria Harbour (https://cd.epic.epd.gov.hk/EPICRIVER/marine/?lang=en). (c) Comparison of Chl-a concentration (mg m^-3^) in the southern Hong Kong waters. The red line indicates data from a monitoring site of *in situ* measurement data, and the dashed line indicates remote sensing data from a monitoring site close to the *in situ* measurement site. Their locations are 22.1917°N, 114.0790°E and 22.18750°N, 114.1042°E. Remote sensing data shows a similar trend with the *in situ* monitoring data.

Here, using an *in situ* measurement dataset collected by EPD and remote sensing dataset collected by MODIS, we apply two tools of EDM, unified information-theoretic causality (UIC) (Osada et al., 2023) to detect causal environmental factors, and Multiview-distance regularized S-map (MDR S-map) (Chang et al., 2021) to quantify interaction strengths of the causal factors, and forecast dynamics of Chl-a in Hong Kong waters. In this study, we first identified causal variables using UIC and quantified the causal effects using MDR S-map.

Then, we conducted near-future forecast of Chl-a dynamics in Hong Kong waters using the time series of Chl-a and causal variables. For *in situ* measurement data, salinity showed the strongest causal effect on Chl-a compared to other environmental factors. However, inputting causal variables into the forecast model (i.e., multi-variable model) did not greatly improve the performance compared to the Chl-a only model (i.e., single-variable model). For remote sensing data, sea surface temperature (SST) showed a significant causal effect on Chl-a at most of the sites and inputting SST improved the performance of the forecast model compared to univariate model using Chl-a only. Lastly, we compared the results of the *in situ* monitoring data and the remote sensing data and discuss the reliability of the remote sensing data for Chl-a monitoring in Hong Kong waters. This study is the first to quantify the causal effect of environmental factors on Chl-a dynamics in Hong Kong coastal waters using EDM, providing perspectives for real-world environmental monitoring, management, and forecasting.

## 2. Materials and methods

### 2.1 In situ measurement data

Monthly time series of Chl-a and environmental factors, including temperature, salinity, pH, turbidity, total nitrogen (TN), and total phosphorus (TP) and silica (as SiO_2_), were collected by Environmental Protection Department (EPD) in 76 sites in Hong Kong waters. Data were downloaded from the EPD website (https://www.epd.gov.hk/epd/english/top.html) as of 2024 January. Each time series started from 2002 August and continued until 2022 July, containing 240 time points in each site. Sampling was carried out onboard a scientific vessel once a month and samples were collected at 1 m below the sea surface. According to EPD, temperature, salinity, pH, and turbidity were measured on site by CTD profiler (SEACAT19+ Conductivity Temperature Depth, Sea-Bird Scientific, US). Chl-a, TN (the sum of Kjeldahl nitrogen, nitrite nitrogen, and nitrate nitrogen), TP and silica were measured in the laboratory as described in Annual Marine Water Quality Reports of EPD (https://www.epd.gov.hk/epd/sites/default/files/epd/english/environmentinhk/water/hkwqrc/files/waterquality/annual-report/marinereport2024.pdf). N/P ratio was defined as TN divided by TP.

### 2.2 Remote sensing data

Remote sensing data of Chl-a and sea surface temperature (SST) were derived from MODIS (Moderate Resolution Imaging Spectroradiometer) onboard the Aqua satellite platform of NASA (National Aeronautics and Space Administration). Data were downloaded from the official website of NASA Ocean Color (https://oceancolor.gsfc.nasa.gov/) as of 2024 January. Data used were monthly Level 3 data at resolution of 4 km. There were 176 sites in total extracted between latitude 22.13°N and 22.58°N, longitude 113.82°E and 114.52°E, among which 101 marine sites had been selected. Each time series started in August 2002 and continued until July 2022, consisting of 240 time points for each site. Missing values of Chl-a concentration were replaced with 0 because missing values usually indicate that the Chl-a concentration was too low.

### 2.3 Unified information-theoretic causality (UIC)

To detect causal effects of environmental factors on Chl-a, unified information-theoretic causality (UIC) was applied (Osada et al., 2023). UIC is a time series-based, nonparametric causality test, which incorporates the advantages of both convergent cross-mapping (CCM; Sugihara et al., 2012), a causality test in EDM, and transfer entropy (TE; Schreiber, 2000), an information-theory based causality test. Here, TE from process *y* to process *x*, *TE_y_*_→*x*_, assesses how much uncertainty in predicting future value of *y* is reduced, given knowledge of past values of *x*. This shares a similar idea with CCM, which regards *y* as a causal effect of *x* if the neighborhood relationship of time series *x* is able to predict time series *y* by nearest neighbor regression in the time-delay embedding. UIC quantifies information flow between variables in format of TE determined by conditional probability (we call this measure TE, but the mathematical definition of *TE_y_*_→*x*_ is different from that of Schreiber; see Osada et al. 2021 for details), which would be comparison of model performance of cross mapping in this case:

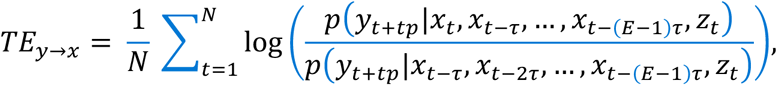

where *x*, *y*, and *z* are a potential effect variable, causal variable, and conditional variable (if available), respectively. In our case, *x*, *y*, and *z* could be Chl-a, environmental factors, and the other potential causal factors, respectively. *p*(*A*|*B*) describes conditional probability: the probability of *A* given *B*. *N* is the length of the library (i.e., number of the vectors in the state space) and *E* is and the optimal embedding dimension when conducting one-step forward forecast in the state space. *t*, *tp*, and *τ* are the time point, time lag between effect variable and causal variable, and time lag of the time series, respectively. To identify the delayed effect of environmental factors on Chl-a, five time-lags, i.e., *tp* = 0, –1, –2, –3, and –4, were tested. These lags suggest that the causal effects occur within the same month, one, two, three, or four months earlier, respectively. For example, when the optimal embedding dimension is five, there is one causal environmental variable, and *tp* = 0, the state vector is represented by the information of an environmental factor of the current month (unlagged), one month ago, two months ago, three months ago, and four months ago, which are used to predict the current state of Chl-a (the numerator in the above equation, which is equivalent to “cross-mapping” in CCM, but is adjusted by the denominator in UIC).

To avoid misidentification of causality caused by seasonality (i.e., synchronized dynamics), a seasonal surrogate test was carried out. For each Chl-a time series, 1000 surrogate time series were generated by computing a seasonal trend (= yearly trend) and shuffling the residuals. If TE from an environmental factor to the original Chl-a time series exceeds TE of over 950 surrogate time series, the environmental factor is regarded as having a significant causal effect on Chl-a (i.e., *p* < 0.05).

For EPD data, causal effects of temperature, salinity, pH, turbidity, N/P ratio, and silica on Chl-a were examined. For MODIS data, causal effects of SST on Chl-a were examined due to the limitation of the availability of environmental variables. To ensure that all variables with different units have the same level of magnitude for comparison and to avoid reconstructing a distorted state space, the time series of all variables were normalized to have a mean of 0 and a standard deviation of 1. This data preprocessing approach differed from that used for our MDR S-map, where the first-differenced time series were normalized (see the following section). Using the first-differenced and normalized time series for UIC showed qualitatively the same results (see Tables S1–S4), but we show the UIC results of the normalized time series in the main text and figures because the interpretation is more straightforward. The computation was conducted using the package “rUIC” (version 0.9.13) (Osada & Ushio, 2021) of R.

### 2.4 Multiview-distance regularized S-map (MDR S-map)

To conduct the near-future forecast of Chl-a dynamics, Multiview-distance regularized S-map (MDR S-map; Chang et al., 2021) was applied. We analyzed the first-differenced and normalized time series to maximize the forecasting accuracy of Chl-a dynamics and to mitigate issues arising from temporal autocorrelation. Before conducting MDR S-map, time series of Chl-a were taken first differenced and normalized and UIC was again conducted to detect causal environmental factors on the differenced Chl-a at each site (Tables S1–S4). Here, a significance test was conducted by a random-shuffled surrogate method (1000 surrogate time series were used to calculate the significance), as the first-differenced time series did not show a clear seasonality. Only sites with significant causal environmental factors were selected for further MDR S-map analysis.

MDR S-map links two existing EDM methods, multiview embedding (Ye & Sugihara, 2016) and regularized S-map (Cenci et al., 2019), which has been proposed to reconstruct large interaction networks when the number of causal variables exceeds the optimal embedding dimension (Chang et al., 2021). The first step of MDR S-map is to determine “multiview distance” describing the “true” neighboring relationship in a high-dimensional state space by ensembling various distances measured in numerous low-dimensional state spaces at the optimal embedding dimension (Ye & Sugihara, 2016). Euclidean distance between every pair of the vectors in the low-dimensional state space, which should be reasonably reliable, is calculated. This procedure is repeated for numerous low-dimensional state spaces, and “ensembled” Euclidean distances among the vectors are calculated by calculating the weighted average among all these distances (the weight is based on the forecast performance of each embedding). The ensembled distance is a good approximation of the “true” distance in the high-dimensional system state.

The second step of MDR S-map is to construct a local linear model, Sequential locally weighted global linear map (S-map, a fundamental tool of EDM; Sugihara, 1994), to fit the time series and make a near-future forecast by including the causal effect from multiple variables,

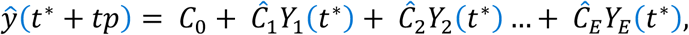

where *t*\*, *E*, and *tp* represent the target time point, the embedding dimension, and forecasting time step, respectively. ŷ*, Y_j_*, and *C_0_* represent the predicted value of *y*, *j*th element of the embedded time series (e.g., lagged Chl-a time series, environmental variables, and so on), and the intercept of the local linear model, respectively. (*Ĉ*_1_, *Ĉ*_2_,…, *Ĉ*_E_) are local linear coefficients. Such Jacobians of the locally approximated linear functions could be defined as the causal effect (or interaction strength). To avoid an overfitting problem when dimension (the number of variables in the linear model) is larger than the time series length, we used regularization (e.g., ridge, lasso, or elastic-net; Cenci et al., 2019) to estimate the coefficients for each time point, *Ĉ* = (*Ĉ*_1_, *Ĉ*_2_,…, *Ĉ*_E_), as follows:

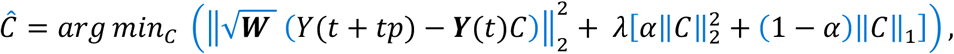

where *C* represents local linear coefficients to be solved, *λ* is the penalized factor set to be selected from 0, 0.001, 0.01, 0.1, 0.5, 1, and 2. *α* is the adjusted parameter set to be 0, balancing the regularization using L1 (||.||_1_) or L2 (||.||_2_) norm of the parameter vector, which means we used the ridge regression. *t* is the time point, and ***W*** is the local weight matrix based on the multiview distances. ***Y***(*t*) = (*Y*_1_(*t*), *Y*_2_(*t*),…, *Y_E_*(*t*)) is a *N* × *E* data matrix (*N* is the number of time points and *E* is the number of nodes, i.e., optical dimension) collecting the time series of all network nodes, and *Y*(*t* + *tp*) = (*Y*(*t*_1_ + *tp*), *Y*(*t*_2_ + *tp*),…, *Y*(*N* + *tp*))*^T^* is *N* × 1 vector representing p-step forward time series data. The solution of the equation depends on parameters *λ* and *α*. All combinations of the parameter values are tested (i.e., grid search) and the best one for each site is determined by normalized mean square error (NMSE) for one-step forward forecast.

In practice, univariate models are defined as models with only Chl-a time series (and its time-lagged values). Under this circumstance, the state space is built by Chl-a and its lagged values. Multivariate models are defined as models with Chl-a and environmental factors (and their time-lagged values). For *in situ* measurement data, multivariate models tested included Chl-a and temperature (“Chl-a + Temp”), Chl-a and salinity (“Chl-a + Sal”), Chl-a and pH (“Chl-a + pH”), Chl-a and turbidity (“Chl-a + Turb”), Chl-a and N/P ratio (“Chl-a + N/P”), and Chl-a and silica (“Chl-a + Sil”). For remote sensing data, the multivariate model tested was built by Chl-a and sea surface temperature (“Chl-a + SST”). Then, the model performance would be evaluated by NMSE. The time series of all variables were normalized to have a mean of 0 and a standard deviation of 1. The computation was conducted using the package “macam” (version 0.1.10) (Ushio, 2025) of R.

### 2.5 Data and code availability

All data used in this study was downloaded from public databases. All scripts and formatted data used in this study are available on Github (https://github.com/sxhuang00/causality_forecast_chl).

## 3. Results

### 3.1 Causal effects of temperature on Chl-a for in situ measurement and remote sensing data

For *in situ* measurement data, several sites in Victoria Harbor showed significant and strong causal effects of water temperature on the Chl-a dynamics (Fig. 2a; *p* < 0.05). Some specific sites in western, southern, and eastern regions and Tolo Harbor also showed significant causal effects of temperature. For remote sensing data, sea surface temperature (SST) exerted significant and common causal effects on Chl-a dynamics in many monitoring sites (Fig. 2b; *p* < 0.05). Some sites in the south and the east that are far from land showed stronger causality of temperature on Chl-a.

**Figure 2.**
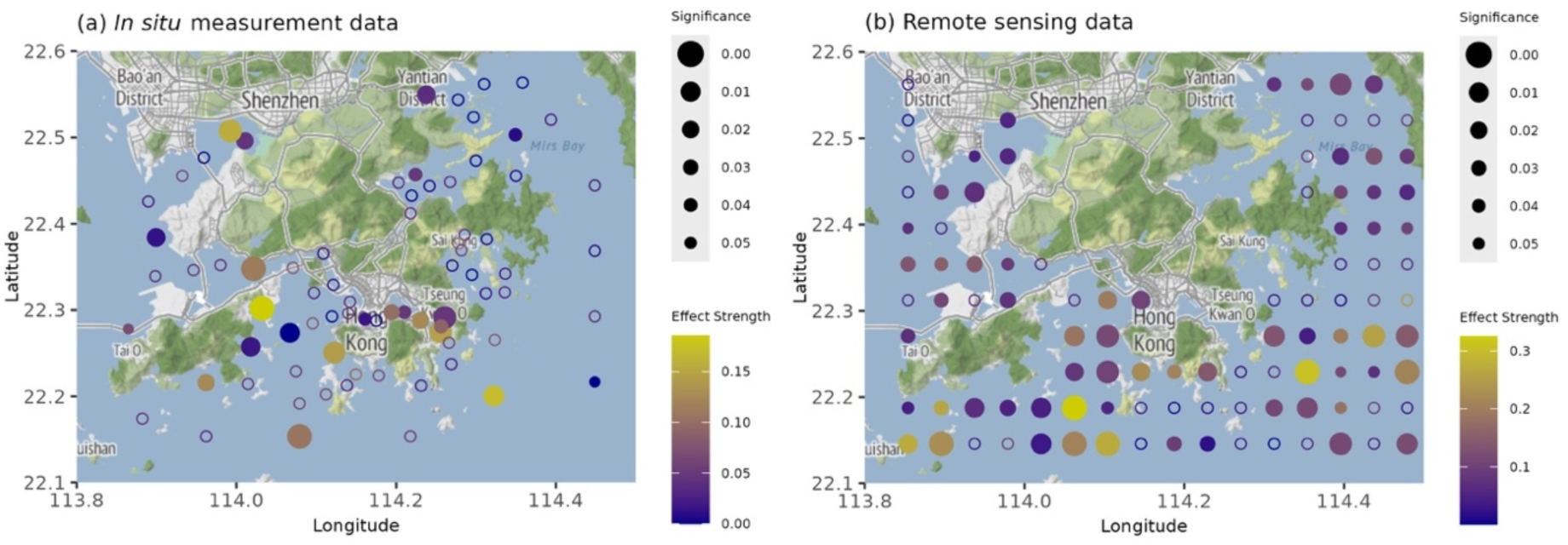
The spatial pattern of causal effect of seawater temperature on Chl-a for (a) *in situ* measurement data (EPD) and (b) remote sensing data (MODIS). Open circles indicate that the causal effect is not significant and filled circles indicate that the causal effect is significant. The circle size indicates the significance of the causal effect with a larger size representing a more significant effect. The circle color indicates the strength of the causal effect (transfer entropy; TE) with lighter color representing a stronger effect of temperature on the Chl-a dynamics.

### 3.2 Causal effects of salinity, pH, turbidity, and nutrients on Chl-a for in situ measurement data

We examined the causal effects of other environmental factors for *in situ* measurement data only, as such data was not available for the remote sensing data. The strengths of the causal effects are summarized in Fig. 3a and the spatial patterns of the causal effects of the environmental factors are shown in Fig. 3b-f.

**Figure 3.**
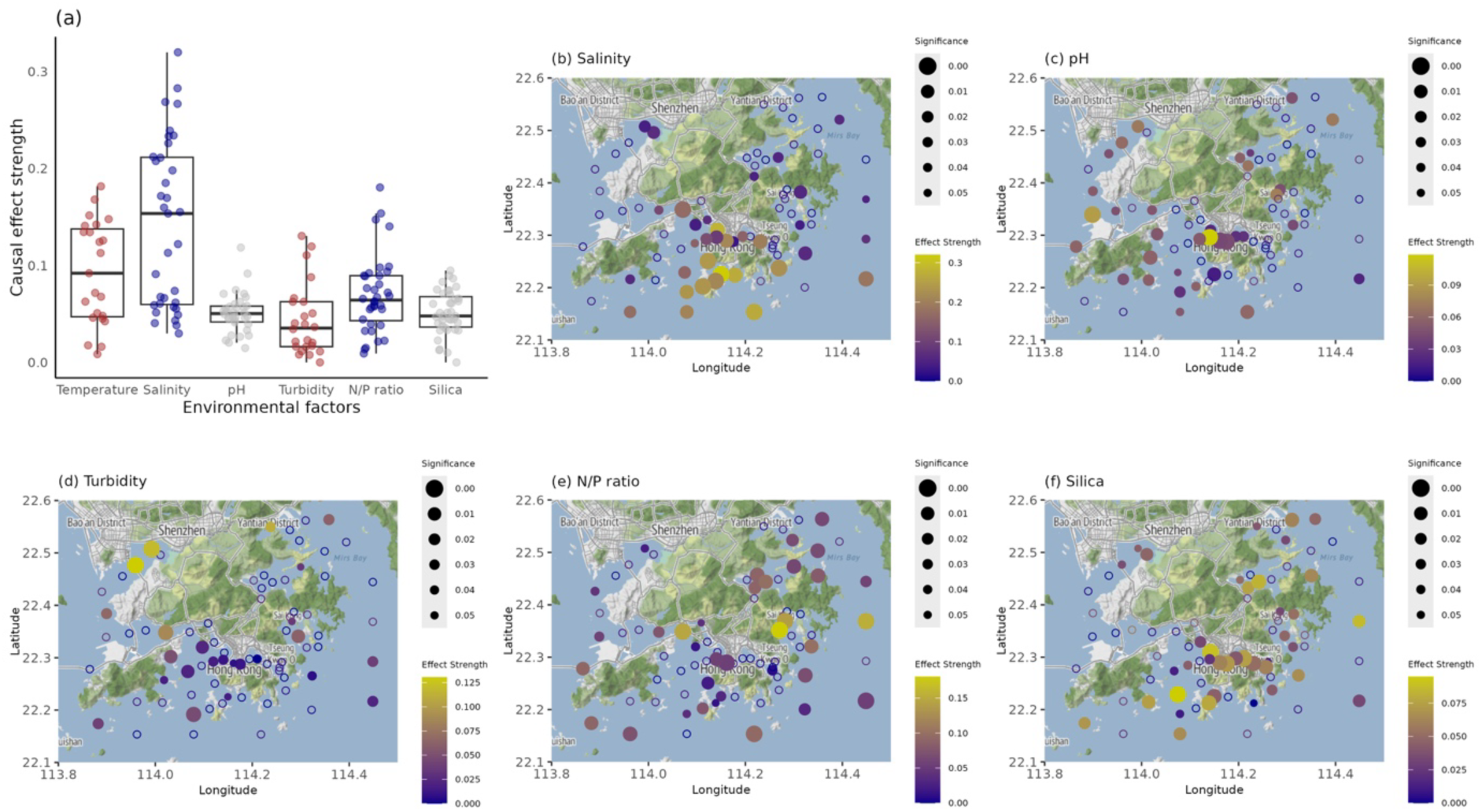
(a) The strength of significant causal effects (transfer entropy, TE) of environmental factors on Chl-a and the spatial pattern of causal effects of environmental factors on chlorophyll-a for *in situ* measurement data: (b) Salinity, (c) pH, (d) Turbidity, (e) N/P ration and (f) silica. Open circle indicates that the causal effect is not significant and filled circle indicates that the causal effect (TE) is significant. The circle size indicates the significance of the causal effect with the larger size representing a more significant effect. The circle color indicates the strength of the causal effect with lighter color representing a stronger effect of temperature on the chlorophyll dynamics.

First, we found that salinity causally influenced the Chl-a dynamics in most monitoring sites in the southern regions and Victoria Harbor (Fig. 3b; *p* < 0.05). The causal effects, measured by TE, of salinity were generally stronger than those of the other environmental factors (average TE = 0.14 for salinity and average TE were 0.09, 0.05, 0.04, 0.07, and 0.05 for temperature, pH, turbidity, N/P ratio, and silica respectively; Fig. 3a). The causal effects of pH were predominantly found in the monitoring sites that were close to shorelines and were located far from oligotrophic regions, such as Mirs Bay in the eastern region (Fig. 3c; *p* < 0.05); however, the causal effects of pH were generally weak (Fig. 3a). The causal effects of turbidity were also generally weak but particularly strong in Deep Bay (Fig. 3d; *p* < 0.05). Causal effects of the N/P ratio were common, especially in Tolo Harbor and Mirs Bay (Fig. 3e; *p* < 0.05). The causal effects of silica showed a similar spatial pattern to those of salinity, particularly in Victoria Harbor and the southern region, but the effect strength was much weaker than that of salinity (Fig. 3f; *p* < 0.05).

### 3.3 Forecasting Chl-a dynamics with univariate and multivariate models using *in situ* measurement data

Using the first-differenced Chl-a time series, we tried to maximize the forecasting accuracy of Chl-a dynamics using MDR S-map. For *in situ* measurement data, multivariate MDR S-map models with different combinations of embedding variables showed similar forecast performance to the univariate model (Fig. 4). Mean NMSEs of “Chl-a”, “Chl-a + Temp”, “Chl-a + Sal”, “Chl-a + pH”, “Chl-a + Tur”, “Chl-a + N/P”, and “Chl-a + Sil” were 0.660, 0.663, 0.666, 0.708, 0.656, 0.708, and 0.690, respectively. Although some sites showed better forecast performance using the multivariate model than using the univariate model, the improvement in the forecast performance when including causal environmental factors was limited.

**Figure 4.**
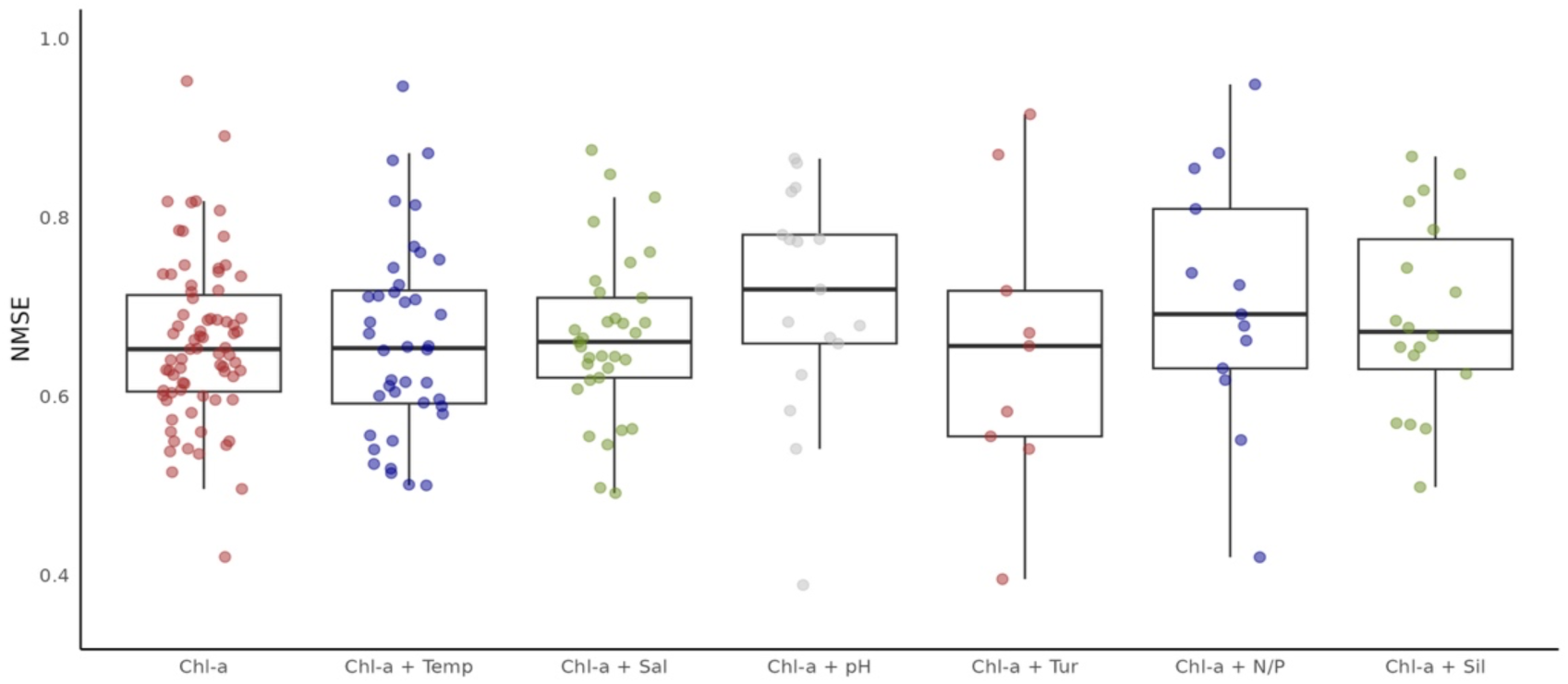
Forecast performance of different MDR S-map models using *in situ* measurement data. The first column indicates the univariate model with differenced Chl-a time series input only. The other columns indicate different combinations of multivariate models with time series of Chl-a and lagged environmental factors input. Each point indicates a monitoring site. Note that the univariate model includes all *in situ* sites, while multivariate models only include sites showing significant causal effect of specific environmental factors on differenced Chl-a, and thus, the number of points for the multivariate models is smaller than that for the univariate model.

### 3.4 Forecasting Chl-a dynamics with univariate and multivariate models using remote sensing data

As for remote sensing data, we analyzed sites showing significant causal effect of SST on differenced Chl-a, and most of them exhibited better forecast performance with the multivariate S-map model than with the univariate S-map (Fig. 5). The mean NMSE of the univariate and multivariate model were 0.632 and 0.574, respectively. Importantly, compared to the univariate model, the multivariate model was better at predicting the high peaks (i.e., Chl-a concentration increases to a high level during algal bloom) and low peaks (i.e., Chl-a concentration decreases to a low level after algal bloom) of Chl-a dynamics without delay or lead (e.g., see Fig. 6).

**Figure 5.**
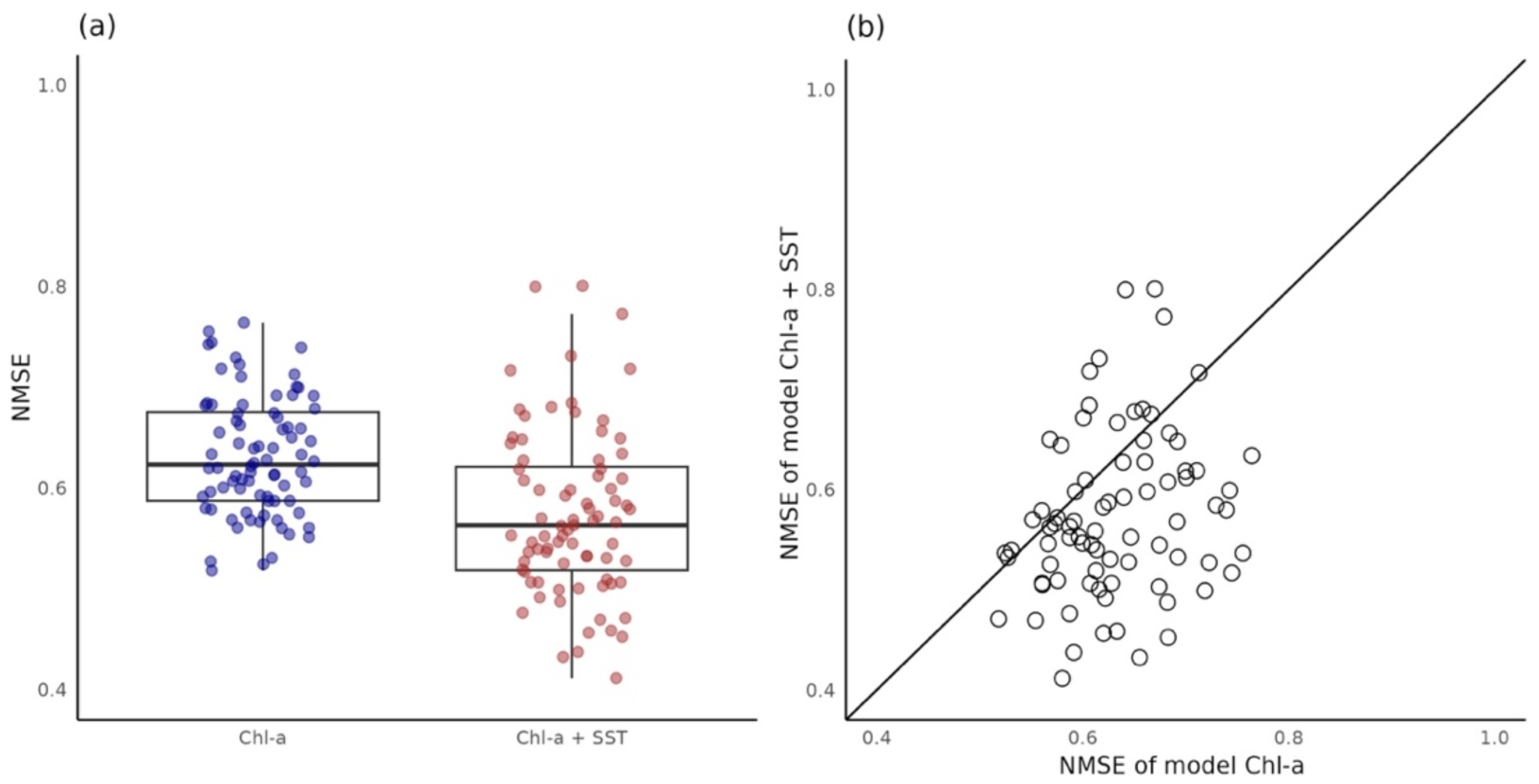
Comparison of forecast performance of different MDR S-map models using remote sensing measurement data. (**a**) Jitter plot of NMSE of univariate model (with Chl-a time series input only) and multivariate model (with time series of Chl-a and lagged SST input. Note that only the sites with significant causal effect of SST on differenced Chl-a are shown). (**b**) Scatterplot of NMSE of univariate model and multivariate model. The solid line indicates the 1:1 line. Points below the 1:1 line indicate the NMSE of multivariate model is smaller than that of univariate model and these sites have better forecast performance using multivariate model.

**Figure 6.**
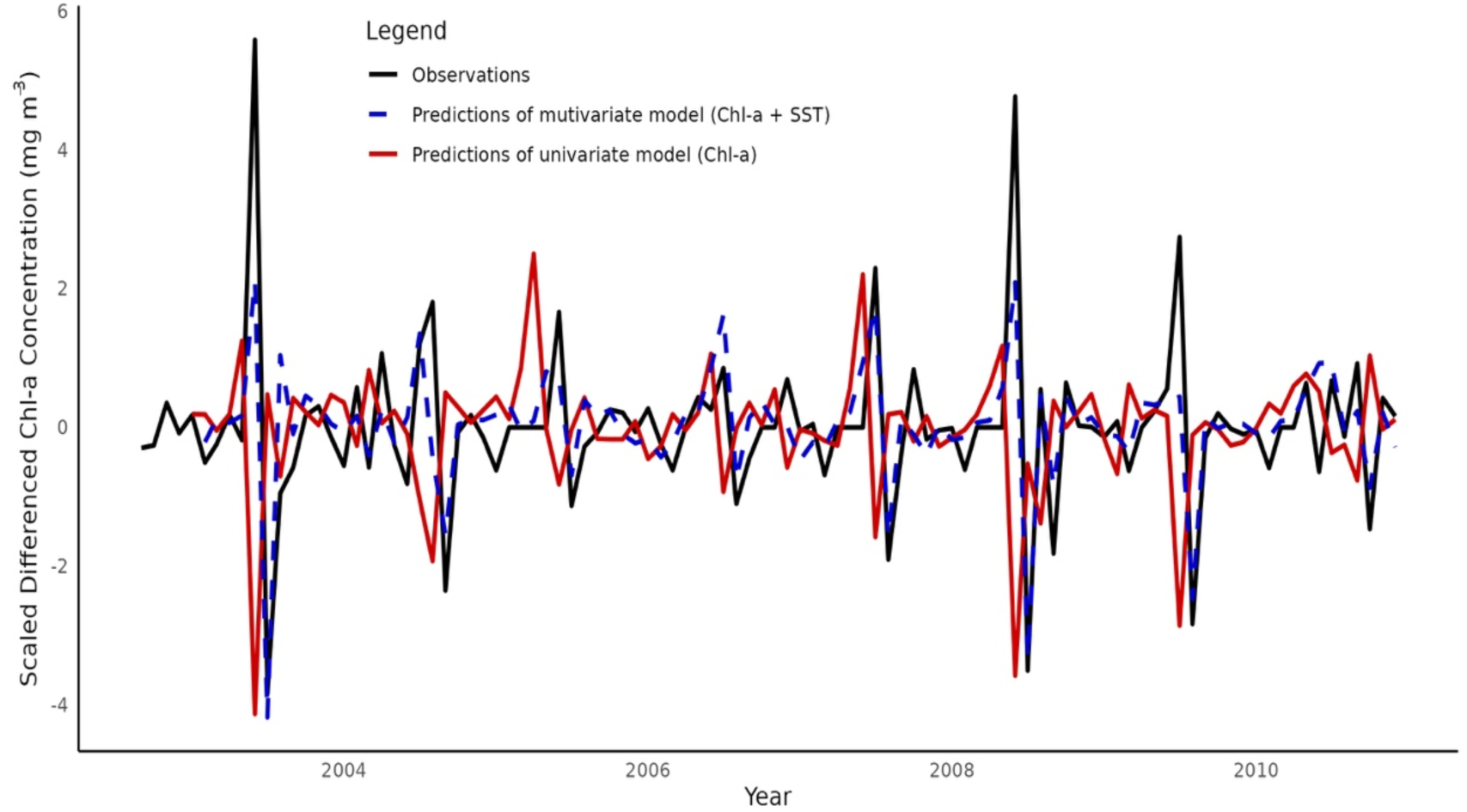
An example of predictions of multivariate model and univariate model of scaled 1^st^ differenced Chl-a concentration time series with NMSE of 0.411 and 0.580, respectively. The original time series belongs to a site located at Port Shelter (22.31250°N, 114.3125°E). To clearly present the data, only the first 100 time points of the time series are displayed here.

## 4. Discussion

In this study, we performed two statistical analyses to gain deeper insights into Chl-a dynamics in Hong Kong waters using Empirical Dynamic Modeling (EDM) tools: (1) identifying causal factors influencing Chl-a dynamics and (2) improving the forecasting accuracy of Chl-a dynamics by incorporating these causal factors. Also, we used two datasets, *in situ* measurements and remote sensing data, to briefly assess the reliability of remote sensing data in a coastal region characterized by highly dynamic nature and high turbidity, which often interfere with data accuracy.

### 4.1 Causal effects of temperature on Chl-a for *in situ* measurement and remote sensing data

First, we found that a causal effect of temperature on Chl-a occurs in some inner corners of semi-closed bays, for example, Deep Bay, Mirs Bay and Tolo Harbor derived from *in situ* measurement data. Temperature dynamics might reflect the occurrences of downwelling inducing weak exchange between these water bodies and open water and long residence time of the bay water (Harrison et al., 2008). Under such circumstances, these water bodies could be considered as an incubator allowing sufficient time for phytoplankton to respond to local nutrient inputs and change the dynamics of Chl-a. Additionally, we found that the strength and spatial patterns of the causal effects of temperature on the Chl-a dynamics were different between *in situ* measurement and remote sensing data (Fig. 2). This discrepancy might be because *in situ* measurement and remote sensing monthly data were collected in different ways. EPD conducted monitoring at irregular intervals, collecting data on different dates in each month. In our analysis, this *in situ* data with irregular intervals was regarded as “regular interval data,” where each data point represents one month. In contrast, monthly remote sensing data was created by averaging daily measurements, potentially providing a more accurate representation of monthly conditions. Thus, while the accuracy of temperature measurement should be higher in the *in situ* data, the constant monitoring intervals of the remote sensing data would be better suited for assessing the impact of temperature on Chl-a dynamics, as EDM requires time series data of consistent intervals.

### 4.2 Causal effects of salinity, pH, turbidity, and nutrients on Chl-a for in situ measurement data

Regarding the other environmental variables, we used the *in situ* data only because of the limited data availability. The causal effects of salinity observed were concentrated in the southern regions and Victoria Harbor (Fig. 3b) in accord with a previous study that showed the establishment of a stable water column by the intrusion of the Pearl River freshwater mass, which could promote algal blooms in these areas (Yin, 2003). The effect of salinity on Chl-a dynamics is stronger than that of other factors (Fig. 3a), suggesting the importance of freshwater discharge and physical processes in algal bloom formation in Hong Kong waters.

pH is related to the availability of inorganic carbon (HCO_3_^−^), which is necessary for photosynthesis, and thus, changes in pH could affect algal growth (Liu et al., 2016). Causal effects of pH are generally found in sites close to the shorelines (Fig. 3c). This might be because pH levels in shorelines are less stable due to more intensive biological activities compared to pH levels in the open ocean area (Duarte et al., 2013).

Primary production of phytoplankton is directly proportional to light availability, which is influenced by turbidity (Domingues et al., 2011). Our results demonstrate that compared to other environmental factors, turbidity generally has a weaker and less common causal effect on Chl-a dynamics (Fig. 3d and a). This might suggest that light availability is a less significant bottom-up factor for phytoplankton dynamics when compared to nutrients, at least in Hong Kong waters. However, in Deep Bay, an estuarine oyster farming zone, turbidity exhibits a notably stronger causal effect. As filter feeders, oysters consume phytoplankton and are more efficient at filtering in clear water (Meeuwig et al., 1998). In oyster farming areas, the impact of turbidity on Chl-a dynamics could be influenced not only by light penetration but also by oyster herbivory, which might make this effect more pronounced in Deep Bay than in other regions.

The causal effect of the N/P ratio was evident not only in eutrophic areas such as Tolo Harbor but also in oligotrophic regions such as Mirs Bay and Port Shelter (Fig. 3e). This indicates that nutrient limitation, which is indicated by the N/P ratio, may be a factor controlling the Chl-a dynamics in these regions. The similar pattern of causal effect of silica with salinity (Fig. 3f and b) could suggest that the availability of silica is affected by fresh river discharge. A previous experimental study demonstrated that available silicon in benthic sediments is subjected to release into the overlying water column for plankton uptake in estuarine and continental shelf environments with lower salinity (Qin & Weng, 2006).

Overall, the causal effects of environmental factors on Chl-a are site-dependent. The hydrodynamic conditions in each area should be taken into consideration when trying to explain the Chl-a dynamics. As the climate in Hong Kong exhibits distinct dry and wet seasons, the effects of rainfall and freshwater discharge (with salinity serving as an indicator) on Chl-a may be temporally dynamic (Lee et al., 2006). Therefore, future studies could focus on how changes in rainfall and freshwater discharge in different seasons control algal bloom. Additionally, water column stability is related to tidal flush, upwelling and downwelling induced by strong wind (Yin, 2003). However, tide current speed, wind speed, and wind direction were not explicitly included in our study sites due to the limitation to accessing such data. Thus, the possible causality of horizontal or vertical movement of water induced by tide or monsoon could not be directly evaluated here. Further study could focus on datasets such as those of Hong Kong Observatory (https://www.hko.gov.hk/en/index.html) for meteorology parameters or Hong Kong Tidal Stream Prediction System (https://current.hydro.gov.hk/main/download.php?lang=en) for oceanographic parameters to draw a fuller picture of algal bloom mechanisms in different water districts.

### 4.3 Forecasting Chl-a dynamics with univariate and multivariate models

We combined the UIC-based causality detection and MDR S-map to conduct near-future forecasting of Chl-a dynamics. For the *in situ* measurement data, multivariate models (i.e., models with Chl-a and other environmental variables) showed similar forecast performance with the univariate model (i.e., Chl-a only model) (Fig. 4). Possible explanations for this result include: 1) The monthly time series data could not effectively capture the fluctuations in nutrient levels and the corresponding responses of Chl-a (or phytoplankton). It has been shown that nutrients delivered by tidal currents and atmospheric inorganic nitrogen can be rapidly consumed by phytoplankton within a timescale of just a few hours (Lo et al., 2025), suggesting that finer temporal resolution data is necessary to make more accurate near-future forecasting. 2) Although the duration of physical processes (e.g., stratification induced by river discharge and downwelling caused by monsoons) usually lasts for a few months, neither salinity nor temperature is a direct indicator of these processes. Incorporating “indirect” indicators of these physical processes might not be sufficient to improve the forecasting accuracy of the univariate model. Future efforts could explore various combinations of factors across different sites at different time scales, considering that the mechanisms behind algal bloom formation may differ in different water bodies.

For the remote sensing data, we found a significant improvement in forecast performance of the multivariate MDR S-map compared to the univariate MDR S-map in most of the sites (Fig. 5), which highlights the potential of EDM that utilizes multiple variables and remote sensing data for monitoring and forecasting Chl-a dynamics. The multivariate model with SST input was also better at capturing the high peaks and low peaks than the univariate model (Fig. 6), suggesting that algal bloom outbreaks are affected by temperature dynamics. The clear improvement in the multivariate model performance for remote sensing data compared to *in situ* measurement data may be attributed to the robustness of monthly average data derived from daily observations, even in the presence of missing values. In contrast, the irregular monitoring interval of time series (EPD conducted sampling on an irregular date of each month) may hinder the performance of EDM.

### 4.4 Comparison of the *in situ* measurement data and remote sensing data, and perspectives for future Chl-a monitoring and forecasting

Although Chl-a dynamics of *in situ* measurement data and remote sensing data at some neighboring sites showed similar trends (Fig. 1c), the causal effect of temperature on Chl-a showed different spatial patterns using these two datasets (Fig. 2). The significant causal effect of temperature was clearly stronger and more common for remote sensing data. Also, the MDR S-map showed improved forecast performance only for remote sensing data. This was expected since EDM is developed for recovering the trajectories of variables coupling with each other and is supposed to demonstrate higher forecast performance if these variables have stronger causal links. Our results reveal that different methods of collecting monthly data can lead to different patterns. Therefore, a consistent sampling interval is recommended to improve forecast performance when applying EDM.

Future efforts should focus on integrating other data resources and analysis methods, including chlorophyll types and/or algae species information and neural network-based algorithms. First, our analysis did not include any functional and/or species information of algae. Different HAB species may have different physiological and population-level characteristics (Chen et al., 2023), and including them in the model could provide better forecasting performance and more detailed information about the algal dynamics (Xi et al., 2021). In addition, an advanced deep-learning model that utilized Chl-a data across a broad spatiotemporal scale in a coastal ocean (Zhang et al., 2025) has provided a potential solution for resolving the missing observation issue of remote sensing data. Further, we could integrate time series data of Chl-a concentrations with historical algal bloom incidents. By employing classifiers such as support vector machines (Keerthi et al., 2001, a machine learning method), we can identify patterns of Chl-a and environmental factors prior to algal bloom outbreaks. This approach could ultimately enhance our ability to predict algal bloom events.

## 5. Conclusions

In the present study, we revealed causal effects of environmental factors on Chl-a and their potential for improving the performance of forecasting Chl-a in Hong Kong waters utilizing a nonlinear time series analysis called Empirical Dynamic Modeling (EDM) and two parallel sets of twenty-year time series data from *in situ* measurements (provided by the Environmental Protection Department; EPD) and remote sensing data (from MODIS). For *in situ* measurement data, salinity exhibited the strongest causal effect on Chl-a compared to other environmental factors, suggesting the importance of oceanographic processes, such as stratification induced by freshwater discharge, in algal bloom formation. As for remote sensing data, SST showed a significant causal effect of Chl-a at most sites and a multivariate model including Chl-a and SST outperformed the univariate model at most sites, highlighting the potential of the multivariate models of EDM. Although the *in situ* measurement data and remote sensing data showed similar Chl-a dynamics in Hong Kong waters, our causal analysis and forecasting model revealed several differences between the two datasets. These findings suggest that accounting for data characteristics (e.g., monitoring intervals) is essential for achieving more efficient and effective monitoring. Overall, this study demonstrates the application of nonlinear time series analysis, EDM, to monthly Chl-a dynamics derived from *in situ* measurements and remote sensing and shows how such approaches can provide insights into Chl-a dynamics in Hong Kong waters. To enhance the water quality monitoring and improve forecasting HAB occurrence in Hong Kong waters, future studies may consider incorporating physical dynamics, developing methods to mitigate the effects of irregular sampling intervals, including species and/or population characteristics, and exploring the potential of other data analysis approaches such as deep learning.

## Data and Code Accessibility

All data used in this study was downloaded from public databases. All scripts and formatted data used in this study are available on Github (https://github.com/sxhuang00/causality_forecast_chl).

## Declaration of generative AI use

We used generative AI tools to polish the English language and improve the clarity of the text. All AI-generated suggestions were manually reviewed and verified for accuracy and clarity.

## Acknowledgments

We thank Takamitsu Ohigashi and Yining Xu for their assistance in data analysis. We thank Mengqiu Wang for her valuable advice on the use of remote sensing data. This research was supported by The Hong Kong University of Science and Technology Startup Fund to MU.

## Author contributions

SH and MU conceived research; SH and MU designed research; SH analyzed the data with help from MU; MU wrote a custom function to perform the MDR S-map; SH and MU wrote the first draft, discussed the results, and completed the manuscript.

